# Sexually Dimorphic DNA Damage Responses and Mutation Avoidance in the Mouse Germline

**DOI:** 10.1101/2020.06.16.155168

**Authors:** Jordana C. Bloom, John C. Schimenti

## Abstract

Germ cells specified during fetal development form the foundation of the mammalian germline. These primordial germ cells (PGCs) undergo rapid proliferation, yet the germline is highly refractory to mutation accumulation compared to somatic cells. Importantly, while the presence of endogenous or exogenous DNA damage has the potential to impact PGCs, there is little known about how these cells respond to stressors. To better understand the DNA damage response (DDR) in these cells, we exposed pregnant mice to ionizing radiation (IR) at specific gestational time points and assessed the DDR in PGCs. Our results show that PGCs prior to sex determination lack a G1 cell cycle checkpoint. Additionally, the response to IR-induced DNA damage differs between female and male PGCs post-sex determination. IR of female PGCs caused uncoupling of germ cell differentiation and meiotic initiation, while male PGCs exhibited repression of piRNA metabolism and transposon de-repression. We also used whole genome single-cell DNA sequencing to reveal that genetic rescue of DNA repair-deficient germ cells (*Fancm*^*-/-*^) leads to increased mutation incidence and biases. Importantly, our work uncovers novel insights into how PGCs exposed to DNA damage can become developmentally defective, leaving only those genetically fit cells to establish the adult germline.

## Introduction

Specification of the germline in humans and mice occurs during embryonic development (Ginsburg et al. 1990), during which PGCs undergo a rapid expansion to populate the fetal gonad (Gomperts et al. 1994). Early in mouse embryogenesis, PGCs are specified as a group of ∼45 cells in the epiblast of 6-6.5 post-fertilization embryos (E6-6.5) (Ewen and Koopman 2010). After specification, PGCs both proliferate and migrate to the location of the future gonads where they undergo roughly 9 population doublings over the span of 7 days to reach a peak population of ∼25,000 cells (Nikolic et al. 2016). These PGCs form the founding germ cell population from which the entire adult germline in both females and males is established (Tam and Snow 1981).

Perturbations to PGC development, especially those that cause accumulation of mutations, can profoundly impact the function and quality of the germline at all subsequent stages of development. In particular, early mutational events in PGCs would be expanded clonally, thus pervading the adult germ cell population. Remarkably, the spontaneous mutation rate in gametes is ∼100 times lower than that of somatic cells (Milholland et al. 2017), suggesting that PGCs, and subsequent stages of gametogenesis, have a highly effective DDR. The ability to suppress mutation transmission in germ cells is essential for maintenance of the germline’s genome integrity, and thus, genetic stability of species and avoidance of birth defects. However, how this suppression is achieved is still incompletely understood.

Studies examining the impact of exogenous genotoxic stressors on germ cells have largely focused on postnatal germ cell development (Russell et al. 1981; Favor 1999; Rinaldi et al. 2017; Enguita-Marruedo et al. 2019; Singh et al. 2018). Fetal germ cells comprise a small population of cells that are difficult to access, but there are some reports in the literature indicating that PGCs are hypersensitive to DNA damage (Hamer and de Rooij 2018). These studies show that mutations in several DNA repair genes impact PGC development, but have subtle effects on other embryonic- and post-natal cell types (Agoulnik et al. 2002; Luo et al. 2014; Luo and Schimenti 2015; Nadler and Braun 2000). Furthermore, germ cells in gastrulating mouse embryos readily undergo apoptosis in response to low dose IR, leading to depletion of the PGC pool; however the underlying mechanisms were not delineated (Heyer et al. 2000). During normal PGC development, some cells are lost through BAX-mediated apoptosis, implying that there are robust quality control mechanisms present and engaged in these germ cells even under physiological conditions (Stallock et al. 2003; Rucker et al. 2000). Exposure to environmental genotoxic agents during fetal development has the potential to impact not only the fetus, but also the future offspring of the fetus through its developing germline. In addition to genetic effects, intrinsic and extrinsic stressors may evoke epigenetic changes to fetal germ cells, possibly impacting fertility and causing adverse health outcomes in subsequent generations. However, our understanding of how PGCs respond to stressors remains under-explored.

DNA damage in the form of double strand breaks (DSBs), which can arise spontaneously (for example during DNA replication) or induced by extrinsic exposures, are particularly dangerous to the genome because they can cause gross chromosomal rearrangements and insertions/deletions (indels). Consequently, the cellular responses to DSBs, commonly induced experimentally by IR, have been studied in many contexts (Ciccia and Elledge 2010; Featherstone and Jackson 1999). While DSBs can be damaging to any cell type in the body, the potential consequences are greater for stem cell populations that would propagate mutations to all progeny cells. Therefore, the DDR in these cells is of particular importance. A tractable system for studying stem cells in mammals are mouse embryonic stem cells (mESCs), and they are highly sensitive to IR exposure compared to other well-studied cell types such as mouse embryonic fibroblasts (MEFs) (Hong and Stambrook 2004; Chuykin et al. 2008; Suvorova et al. 2016; Tichy and Stambrook 2008). Interestingly, PGCs have several properties resembling mESCs, including rapid proliferation, low mutation rate, and similar transcriptomes (Hong et al. 2007; Cervantes et al. 2002; Grskovic et al. 2007). Under proper culture conditions, mESCs can even be differentiated into PGC-like cells (PGCLCs) in just a few days (Hayashi et al. 2011). This raises the possibility that PGCs and mESCs have similar DDRs that are distinct from terminally differentiated cells, and that ESCs, which are very easily cultured, can be used as a guide for studying DDRs of other stem cells including PGCs.

In this study, we examined the PGC response to IR-induced DNA damage at two distinct stages of development: 1) when PGCs are bipotent prior to sex determination at E11.5, and 2) subsequent to the initiation of sex determination at E13.5 (Endo et al. 2019). At E11.5, female and male fetal gonads are morphologically indistinguishable from one another, but by E13.5 the gonads are morphologically distinct (Koubova et al. 2006). Additionally, the developmental trajectories of male and female PGCs begin to diverge at this time. Female germ cells begin to undergo meiotic initiation at E13.5, while male germ cells continue on a mitotic cell cycle program before becoming quiescent ∼2 days later (Anderson et al. 2008). By examining the IR-induced DDR in PGCs both before and after sex determination, we uncovered novel developmental context-dependent responses to DNA damage. We show that before sex determination, irradiated PGCs lack a G1 cell cycle checkpoint similar to mESCs. After sex determination, we show that male PGCs re-gain G1 checkpoint activity while female PGCs do not, and instead prematurely initiate an abortive oogonial differentiation program. We also assessed mutational burden and the role of the G1 cell cycle checkpoint in mutation prevention in an intrinsic DNA damage model, *Fancm*-deficient mice, that exhibit p21-dependent PGC depletion in males (Luo et al. 2014). Overall, our studies reveal the importance of the G1 checkpoint in preventing accumulation of complex mutations in the germline, and the differentiation of the DDR during germ cell development.

## Results

### Mouse PGCs Lack a G1 Cell Cycle Checkpoint

In a proliferating population of cells, acute DNA damage can activate cell cycle checkpoints at a number of different cell cycle stages (Shaltiel et al. 2015). These checkpoints give cells a chance to respond to the damage and can lead to a shift in the population’s cell cycle distribution compared to control, undamaged cells. The canonical DDR in many cell types involves activation of a checkpoint at the G1 stage of the cell cycle, but notably, this checkpoint is absent in mouse and primate ESCs (Hong et al. 2007; Hong and Stambrook 2004; Fluckiger et al. 2006). The speculated reason for this strategy is that rather than attempting repair of a mutational load sufficient to stop the cell cycle at G1 as do most somatic cells, ESCs sustaining substantial DNA damage of a cell-deleterious nature get culled subsequently by other mechanisms. Mouse neural stem and progenitor cells (NSPCs) and hematopoietic stem cells (HSCs) also do not activate a G1 cell cycle block in response to IR (Roque et al. 2012; Brown et al. 2015).

To determine whether PGCs lack a G1 DNA damage checkpoint, we exposed pregnant mice at E11.5 to 5 Gy IR, then 8 hours later dissected fetal gonads for flow cytometric cell cycle analysis. The mice expressed GFP in germ cells (Szabo et al. 2002), enabling us to distinguish PGCs from gonadal somatic cells. The IR treatment caused a marked shift in the PGC cell cycle distribution, indicative of an absent G1 cell cycle checkpoint (Figure 1A). This response was similar to that previously reported for mESCs (Hong and Stambrook 2004), an observation we confirmed here (Figure S1). The dramatic decrease of PGCs in G1 in turn altered the distribution of cells in S- and G2/M-stages (Figure 1B-D) indicating the presence of a robust G2 checkpoint arrest similarly identified in mESCs.

**Figure 1.**
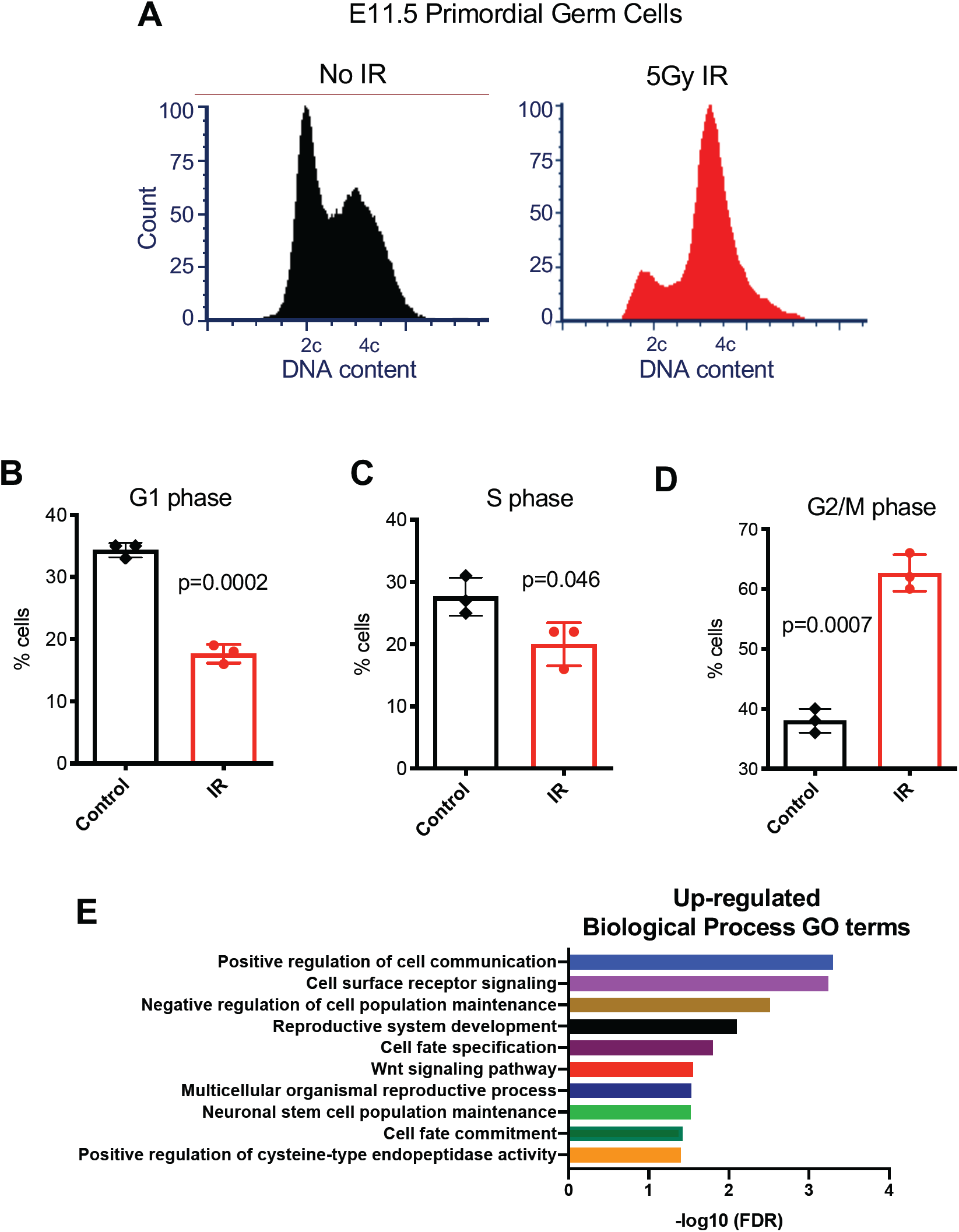
PGCs at E11.5 lack a G1 cell cycle checkpoint in response to irradiation-induced DNA damage. A. Representative cell cycle profiles of control and treated primordial germ cells. B. Percentage of G1 phase cells in both control and treated conditions. C. Percentage of S phase cells in both control and treated conditions. D. Percentage of G2/M phase cells in both control and treated conditions. E. Significantly enriched Gene Ontology terms among up-regulated genes in response to irradiation at E11.5.

### Irradiation Leads to Decreased piRNA Metabolism and Transposon de-repression in the Fetal Male Germline

To better understand the molecular nature of the DDR in bipotent PGCs, we performed RNA-seq on PGCs purified from irradiated embryos compared to control PGCs at the same time point in development (E11.5). Comparisons between irradiated and unirradiated samples 4 hours after treatment revealed 282 differentially expressed genes with 124 genes up-regulated two-fold or higher and 18 genes down-regulated two-fold or lower (Table S6). Gene Ontology analysis (Mi et al. 2019) highlighted a number of significant terms among the genes up-regulated in response to IR (Figure 1E), most notably cell-to-cell communication, factors involved in stem cell population maintenance, and activation of apoptotic processes through cysteine endopeptidase activity (Earnshaw et al. 1999).

Having found that E11.5 PGCs lack a G1 cell cycle checkpoint, we next asked if and when the G1 checkpoint response is re-established in this lineage. Establishment of a G1 checkpoint response would indicate that the lineage re-gained a more canonical DDR, typical of most differentiated cells. Strikingly, irradiation of male embryonic germ cells at E13.5 induced G1 cell cycle arrest, indicating that they acquired this ability for the first time in their development (Figure 2A-D). To gain insight into the molecular consequences of DNA damage in these cells, we examined the transcriptomes of irradiated E13.5 male PGCs to unirradiated controls using RNA-seq. During normal male PGC development, there is widespread demethylation of the genome (Ernst et al. 2017). This global DNA demethylation leads to de-repression of silenced transposons, but also coincides with an upregulation of piRNA signaling to repress transposon expression (Rojas-Ríos and Simonelig 2018). Activation of piRNA signaling is crucial for normal developmental progression of fetal male germ cells (Nguyen and Laird 2019). Gene Ontology (GO) analysis highlighted a number of gene categories down-regulated in response to IR, revealing perturbations to critical male germ cell-specific developmental pathways related to gene silencing, cell differentiation and piRNA signaling (Figure 3A).

**Figure 2.**
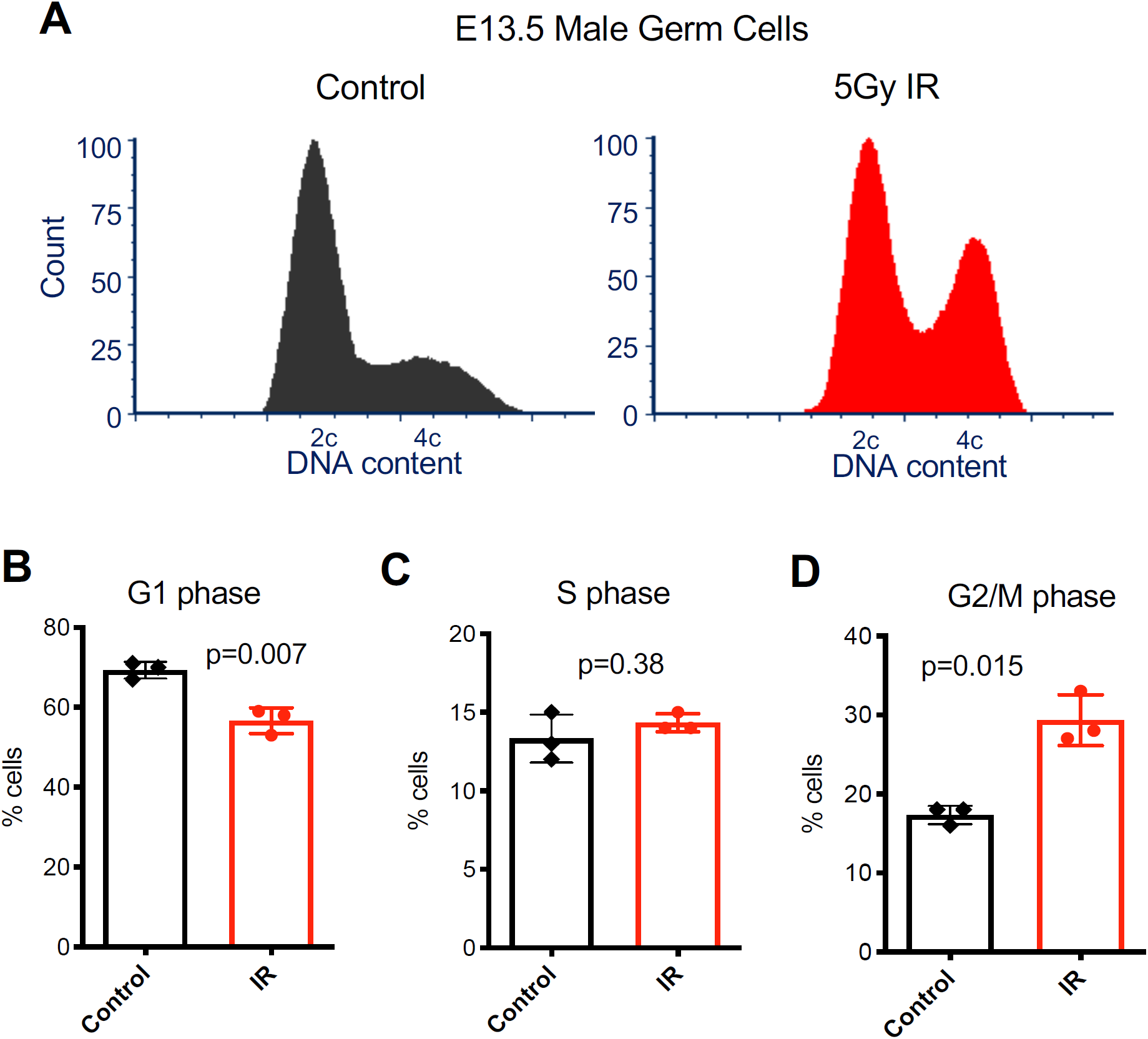
E13.5 male germ cells acquire a DNA damage responsive G1 cell cycle checkpoint. A. Representative cell cycle profiles of control and treated E13.5 male germ cells. B. Percentage of G1 phase cells in both control and treated conditions. C. Percentage of S phase cells in both control and treated conditions. D. Percentage of G2/M phase cells in both control and treated conditions.

**Figure 3.**
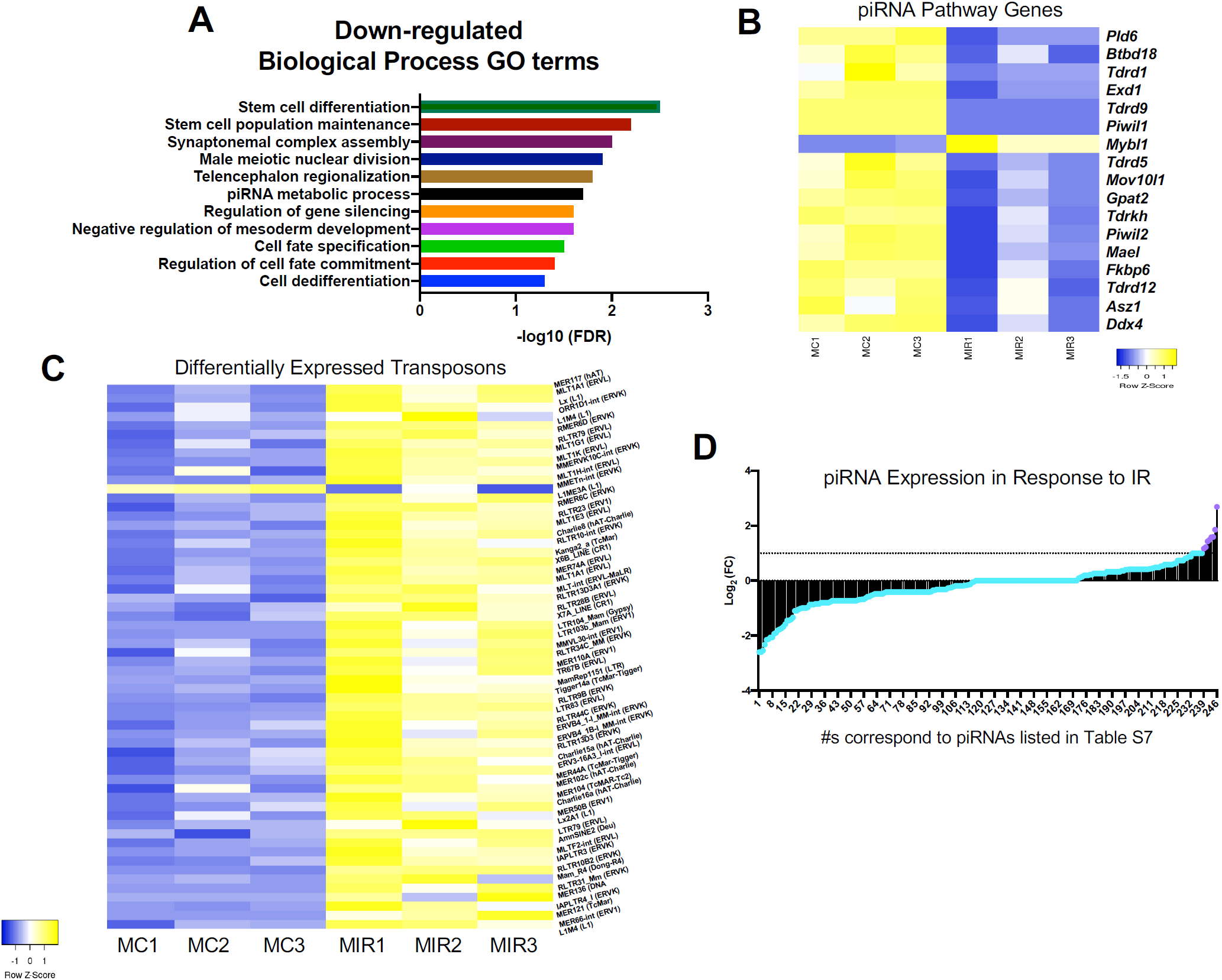
Irradiation exposure at E13.5 leads to down-regulation of piRNA metabolism and de-repression of transposons in male germ cells. A. Significantly enriched GO terms among down-regulated genes in response to irradiation in E13.5 male germ cells. B. Expression of piRNA metabolic process genes in control and irradiated E13.5 male germ cells. C. Heatmap of differentially expressed transposons between control and irradiated samples with an adjusted p-value<0.05. D. Fold change in piRNA expression in response to IR from small-RNA sequencing. The majority of the piRNAs (shown in blue) are not significantly up-regulated in response to IR. (MC=male no IR control; MIR= male IR treated)

To further understand the DDR in E13.5 male germ cells, we examined the expression of genes associated with piRNA metabolism in our dataset. The vast majority (all except one) were robustly down-regulated in response to IR (Figure 3B). Next, we asked whether the down-regulation of piRNA metabolism led to transposon activation in the irradiated male germ cells. We observed that the majority of expressed transposons fall into families that are capable of transposition (Figure S2) (Deniz et al. 2019). Additionally, when we compare the expression of all differentially expressed transposons with an adjusted p value<0.05, all but one were de-repressed in irradiated germ cells (Figure 3C).

Finally, we conducted small RNA sequencing to examine the expression of piRNAs and observed that the majority were down-regulated in response to IR (Figure 3D). Therefore, induction of DNA damage in mouse embryonic germ cells suppresses piRNA activity by downregulating both piRNA biogenesis factors and the piRNAs themselves.

### Irradiation of Female Germ Cells Leads to an Uncoupling of Oocyte Differentiation and Meiosis

As described above, post-sex determination male PGCs (E13.5) acquire a G1 checkpoint response as they come to the end of their highly proliferative phase. With respect to female PGCs, like their counterparts at E11.5, IR caused a marked shift in the number of cells with 4C DNA content, indicating that they also lacked a G1 checkpoint at this stage (Figure 4A-D). However, the fate of normal female germ cells differs at E13.5, as they begin transitioning into a meiotically competent cell cycle program. Interestingly, the IR-induced cell cycle profile of E13.5 female germ cells did not differ from the profile at E11.5 (Figure 4A). We hypothesized that rather than representing a lack of a mitotic G1 checkpoint, these cells might actually represent prematurely-arising oocytes resulting from IR induction. Consistent with this possibility, we noted that this cell cycle profile was remarkably similar to that reported for normal E14.5 female germ cells (Miles et al. 2010).

**Figure 4.**
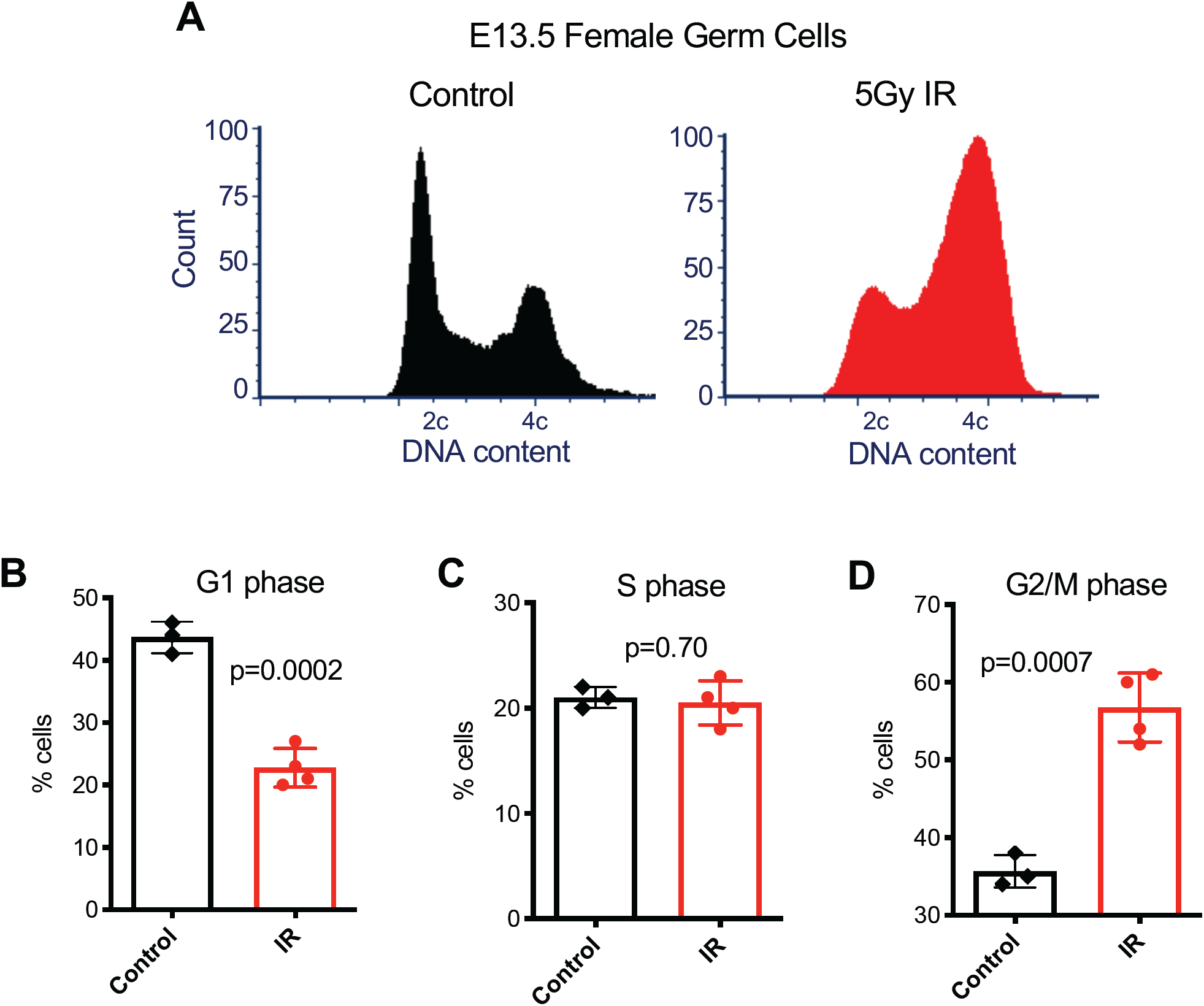
Enrichment of G2/M phase cells in IR exposed E13.5 female germ cells. A. Representative cell cycle profiles of control and treated E13.5 female germ cells. B. Percentage of G1 phase cells in both control and treated conditions. C. Percentage of S phase cells in both control and treated conditions. D. Percentage of G2/M phase cells in both control and treated conditions.

To test this hypothesis, we performed RNA-seq of the IR-exposed E13.5 germ cells, and compared the up- and down-regulated genes to a list of pluripotency-related genes expressed in fetal germ cells (Sangrithi et al. 2017; Lesch et al. 2013). Several genes (23) associated with pluripotency were downregulated in response to IR (Figure 5A), consistent with IR exposure causing premature differentiation of female PGCs. Additionally, comparisons between irradiated and unirradiated E13.5 female germ cells 8 hours after treatment also revealed an increase in retinoic acid (RA) responsive genes (Figure 5B).

**Figure 5.**
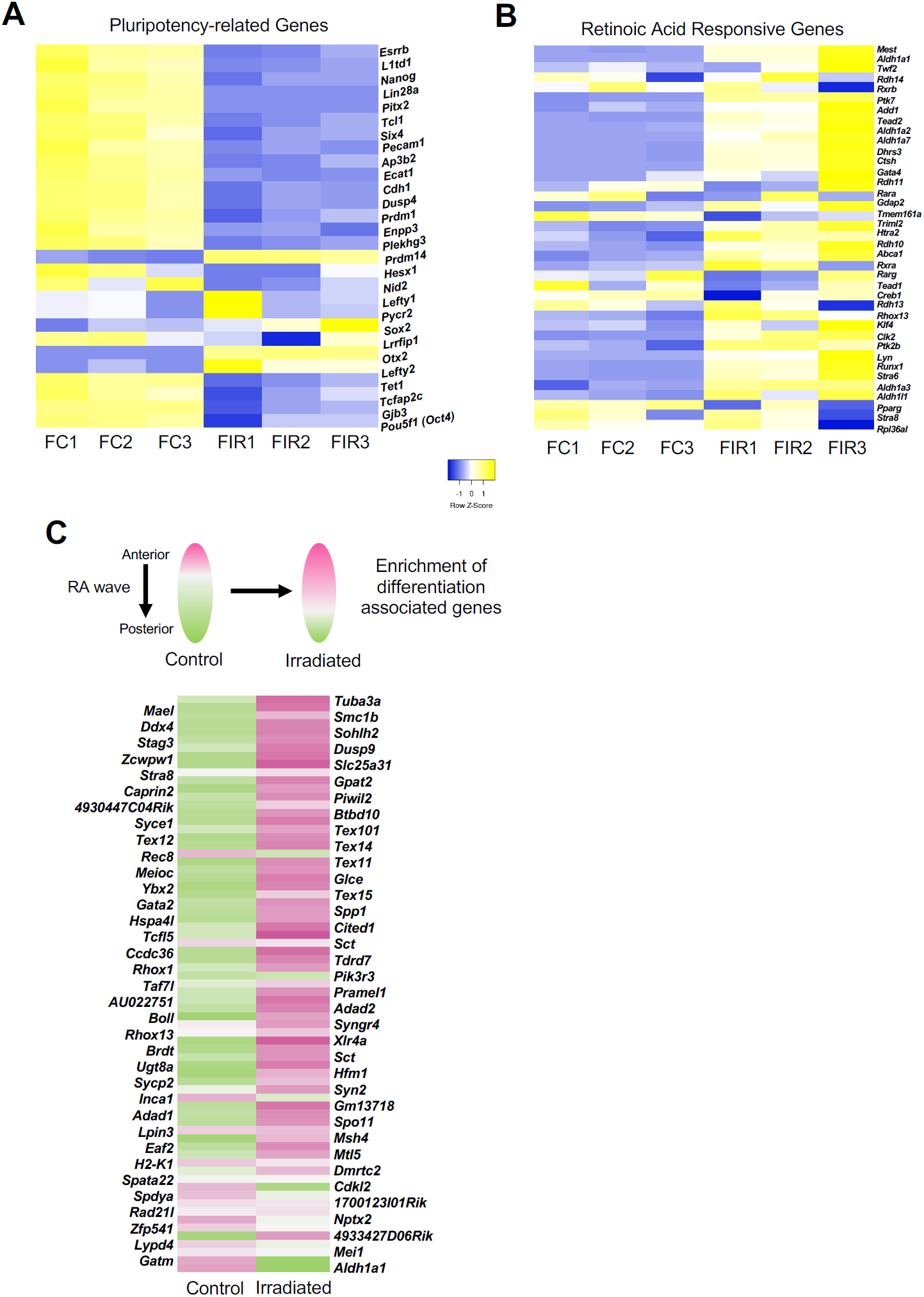
Irradiation exposure at E13.5 leads to pre-mature retinoic acid signaling and meiotic entry disruption in female germ cells. A. Expression of pluripotency-associated genes in control and irradiated E13.5 female germ cells. B. Expression of retinoic acid responsive genes in control and irradiated E13.5 female germ cells. C. Expression of genes associated with spatial development of the fetal ovary in control and irradiated samples. The heatmap is comprised of genes upregulated in response to RA exposure, pink indicates higher expression, green indicates lower expression (genes listed on the left are associated with even rows of the heatmap and genes listed on the right are associated with the odd rows; reference dataset used for comparison from (Soh et al. 2015). (abbreviations: FC=female no IR control; FIR= female IR treated)

Entry into meiosis requires the completion of pre-meiotic DNA replication and an extended Prophase I stage (Speed 1982; Soh et al. 2017). Moreover, initiation of the meiotic program in female germ cells occurs in an anterior to posterior wave of RA signaling in the fetal gonad (Koubova et al. 2006). Therefore, based on the increased expression of RA responsive genes and the dramatic upsurge in G2/M phase cells (Figure 4D), we hypothesized that IR stimulates RA-associated gene expression, which in turn stimulates meiotic entry. To explore this possibility, we took advantage of a published dataset where the embryonic ovary was dissected into thirds and RNA-sequencing performed on the anterior and posterior portions (Soh et al. 2015). Comparison to that dataset revealed that gene expression in irradiated fetal germ cells is more similar to the portion of the embryonic gonad which has been exposed to RA and initiated meiotic entry (Figure 5C).

Upon entry into meiosis, a set of primarily meiosis-specific genes become highly expressed (Sangrithi et al. 2017; Lesch et al. 2013). Surprisingly, analysis of these genes in our dataset revealed that rather than being more highly expressed in germ cells exposed to IR, they are down-regulated (Figure S3). This result, which seemingly conflicts with the IR-induced enrichment of gene expression associated with RA signaling, led us to conclude that IR causes an uncoupling of oogonial differentiation from meiotic entry. Previous work in mouse has demonstrated that oogonial differentiation and meiosis are dissociable from one another (Dokshin et al. 2013), but, importantly, our work has uncovered a mechanism by which DNA damage triggers this dissociation in an otherwise wild-type context.

Why does IR-induced stimulation of RA signaling not lead to premature meiotic entry in E13.5 female germ cells? One possibility relates to one of the master regulators of meiotic entry, *Stra8* (Koubova et al. 2014). *Stra8* stands for “Stimulated by Retinoic Acid 8” and as the gene name implies, its activity is dependent on RA (Koubova et al. 2006). While RA activates transcription of *Stra8*, expression of *Stra8* leads to the formation of a negative feedback loop and self-repression (Soh et al. 2015). This mechanism ensures that meiotic entry initiates only once. We hypothesized that IR-induced DNA damage stimulates *Stra8* prematurely, leading to an inappropriate and irreversible repression of meiosis. If this is indeed the case, then we would predict that the irradiated germ cell samples would have a similar gene expression profile to *Stra8*-deficient female germ cells. Using an RNA-sequencing dataset of female *Stra8* mutant germ cells (Soh et al. 2015), we observed that genes highly expressed in *Stra8* mutants are indeed similarly enriched in the irradiated germ cell samples (Figure S4).

### Rescue of *Fancm*-deficient Germ Cells by Checkpoint Ablation Leads to an Enrichment of Complex Mutations

To assess how germline mutational burden is impacted when DNA damage checkpoints are abrogated, we sought to examine mutation incidence in a PGC proliferation-defective mouse model. We chose to examine *Fancm*-deficient mice in which male, but not female, germ cell reduction could be partially rescued by deletion of the cyclin-dependent kinase inhibitor, *p21* (Luo et al. 2014). This sexual dimorphism in *p21-*mediated germ cell rescue is consistent with our cell cycle results demonstrating a male-specific establishment of the G1 checkpoint in PGCs after sex determination.

*Fancm* is the largest subunit of the Fanconi Anemia core complex, which is named after a chromosomal instability syndrome that leads to cancer predisposition, bone marrow failure, congenital abnormalities and infertility (Joenje and Patel 2001). Studies in cell culture systems have shown that *Fancm* facilitates cell cycle checkpoint activation at sites of arrested DNA replication forks, particularly in the contexts of interstrand crosslinks (ICL) (Deans and West 2009). *Fancm* has also been reported to mediate fork reversal when the lagging strand template is partially single-stranded and bound by the single-stranded DNA binding protein RPA (Gari et al. 2008). Additionally, single molecule studies have indicated that *Fancm* may even be capable of mediating transversal of ICLs by replication forks (Huang et al. 2013).

Using CRISPR/Cas9 genome editing, we simultaneously generated *Fancm* and *p21* null mutations on an isogenic strain background. Characterization of the *Fancm* mutant revealed phenotypic similarities to the previously published mutant including a partial, but significant, rescue of male germ cells in *Fancm*^*-/-*^ *p21*^*-/-*^ double mutants (Luo et al. 2014) (Figure S5). To examine whether rescuing germ cell quantity through checkpoint bypass led to a decrease in germline genome quality, we compared germline mutation incidence between the single- and double-mutants by collecting spermatids from wild-type, *p21*^*-/-*^, *Fancm*^*-/-*^, and *Fancm*^*-/-*^ *p21*^*-/-*^ animals and performing whole genome single-cell DNA sequencing on them. An increase in mutation incidence in the double mutants would indicate that removing the *p21*-mediated G1 checkpoint has a negative impact on germline genome integrity. A mutational burden similar to controls would suggest that *p21* loss facilitates germ cell rescue without impacting germ cell quality. Either outcome will lead to important insights regarding the relationship between cell cycle checkpoint response and germline genome maintenance.

Initial analysis of all detected variants in the cells revealed an over-representation of mutations in double mutant germ cells (Figure 6A). Further analysis comparing the types of mutations per genotype indicated that the point mutation frequency, while enriched in *Fancm*^*-/-*^ *p21*^*-/-*^ double mutant spermatids did not reach statistical significance when compared to other genotypes (Figure 6B). Breakdown of point mutation type (Figure S6) also did not show distinctions between any of the genotypes.

**Figure 6.**
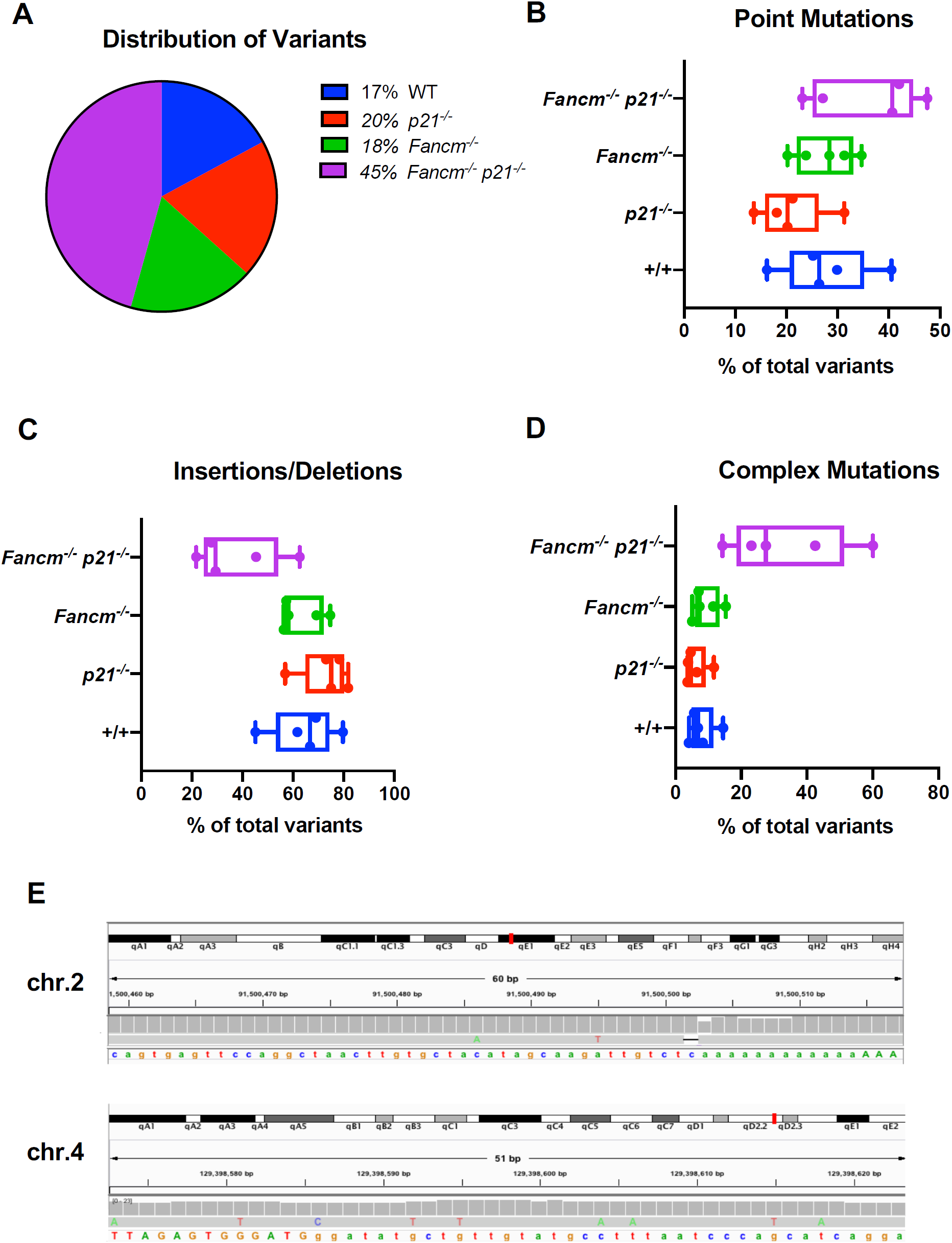
Variant detection in genetically rescued germ cells reveals an alteration mutational profile compared to controls. A. Distribution of total variants with respect to genotype. B. Point mutation frequency shown as a percentage of the total number of variants identified in each cell. (Kruskal-Wallis test, q=0.09 between *p21*^*-/-*^ and *Fancm*^*-/-*^ *p21*^*-/-*^). C. Insertion and deletion (InDel) mutation frequency shown as a percentage of the total number of variants identified in each cell. (Kruskal-Wallis test, q=0.02 between *p21*^*-/-*^ and *Fancm*^*-/-*^ *p21*^*-/-*^) D. Complex variant frequency shown as a percentage of the total number of variants identified in each cell. (Kruskal-Wallis test, q=0.04 between wild-type and *Fancm*^*-/-*^ *p21*^*-/-*^). E. Examples of two complex mutations from *Fancm*^*-/-*^ *p21*^*-/-*^ spermatids. Shown are IGV screen shots.

Next, we examined insertion/deletion (InDel) frequency and, while depleted in *Fancm*^*-/-*^ *p21*^*-/-*^ mutant cells compared to the other genotypes, the differences only reached the threshold for statistical significance when compared to that of *p21*^*-/-*^ cells (Figure 6C). Furthermore, predicted frameshift-causing variants (Figure S7) did not differ statistically between double mutant cells and those from wild-type or single mutants. Notably, clusters of mutations in close proximity to one another were enriched in *Fancm*^*-/-*^ *p21*^*-/-*^ spermatids compared to wild-type (Figure 6D). These complex mutation events were often comprised of InDels along with one or more base substitutions. We defined variants as “complex” if two or more mutations were within 200 nucleotides of one another. Examples of these clustered mutations from individual *Fancm*^*-/-*^ *p21*^*-/-*^ germ cells are shown in Figure 6E. The enrichment of this mutation cluster signature in *Fancm*^*-/-*^ *p21*^*-/-*^ germ cells indicates that the p21-dependent cell cycle checkpoint is important for suppressing propagation of germ cells bearing high levels of complex mutations, or which are experiencing a high number of defective replication forks that would lead to such mutational events.

## Discussion

In this study, we examined DNA damage checkpoint activity in mouse PGCs and identified developmental context-dependent responses before and after sex determination in these cells. We found similarities between the DDR of mESCs (Hong and Stambrook 2004), NSPCs (Roque et al. 2012), HSCs (Brown et al. 2015) and E11.5 PGCs. These similarities have biological and experimental implications. Regarding the former, the results suggest that key features of the DDR are similar amongst distinct stem cell types, from highly pluripotent cells (ESCs) to those dedicated to different lineages (PGCs, NSPCs and HSCs). Experimentally, the data suggest that mESCs, which can be cultured indefinitely and are easily manipulated genetically, could serve as a model for DNA damage responses in PGCs and possibly other cell types that are not easily cultured.

Elucidation of the IR-induced DDR in PGCs following sex determination highlights how perturbations to cell differentiation can be used as a highly sensitive response to preserve the genomic integrity of the final surviving germ cell pool. De-repression of transposons and down-regulation of piRNA signaling in the male germline illustrates the deleterious downstream consequences of DNA damage beyond the direct consequences of the IR itself. Our findings are consistent with a study which showed that in wild-type E13.5 male PGCs, there is a sub-population of cells exhibiting co-occurrence of piRNA pathway down-regulation and up-regulation of apoptotic pathway-associated genes (Nguyen and Laird 2019).

Evidence for RA pathway stimulation in response to DNA damage has also been found in ESCs. In this system, DNA damage induced RA pathway activation promoted cell differentiation through *Stra6. Stra6*, like *Stra8*, is a retinoic acid-responsive gene (Carrera et al. 2013; Serio et al. 2019). We also observed a DNA damage-induced increase in RA signaling and differentiation in female PGCs, but based on the specifics of RA-induced meiotic entry, cannot exclude the possibility that there is a selective enrichment of more differentiated cells among the surviving irradiated cells. This alternative possibility implies that differentiated female PGCs are more resistant to DNA damage than their less differentiated RA-naïve counterparts. It is possible that our timing of exogenous DNA damage with cells primed to tolerate hundreds of programmed meiotic DSBs may distinguish these cells from their less differentiated counterparts, but, importantly, we show that the irradiated cells exhibit inappropriate expression of meiosis-associated genes. Therefore, these germ cells, regardless of whether they represent RA-exposed cells prior to IR or a population induced to differentiate, are not developmentally competent to establish the germline.

Finally, we assessed the quality of individual haploid germ cells in a replication defective mutant. With whole genome single cell DNA sequencing, we were able to highlight the potential ramifications of manipulating DNA damage checkpoints to facilitate increased germ cell survival. Our results indicate that increasing cell survival in a model of germline DNA repair deficiency leads to germ cells with an increased mutational burden. Repair of *Fancm*-associated DNA lesions in other cell types has been shown to involve exposure of single-stranded DNA and repair by low fidelity translesion synthesis (TLS) DNA polymerases (Grompe and D’Andrea 2001). Specifically in PGCs, *Rev7* (a subunit of the TLS DNA polymerase ζ) has been shown to be essential for cell survival with complete loss of PGCs by E13.5 in *Rev7-*deficient mutants (Watanabe et al. 2013). Taken together, these findings provide additional independent support for the idea that enriched variants identified in the double mutant spermatids likely arose from TLS events (Harfe and Jinks-Robertson 2000; Stone et al. 2012).

Notably, there are many genes which lead to germ cell depletion, some via apoptosis and some by slowing DNA replication. In this specific genetic context, the absence of *p21* enables the increased survival of cells with complex mutations via checkpoint bypass. It will be important to continue exploring the impact of these various contexts on *de novo* germline variation throughout germ cell development. Overall, our findings provide novel insight into how the germline minimizes mutation transmission to future generations when exposed to DNA damage and replication stress *in utero*.

## Materials and Methods

### Mouse Models

The use of mice in this study was approved by Cornell’s Institutional Animal Care and Use Committee. B6;CBA-Tg(Pou5f1-EGFP)2Mnn/J transgenic mouse strain, commonly referred to as Oct4deltaPE-GFP, (Jackson Laboratory stock # 004654) were used to purify PGCs. For the irradiation experiments, mice were placed in a ^137^cesium irradiator with a rotating turntable and exposed to the dose of radiation specified.

### Generation of *Fancm* ^*em1/Jcs*^ and *p21* ^*em1/Jcs*^ Mice

*p21*^*em1/Jcs*^ was generated using CRISPR/Cas9-mediated genome editing. The sgRNA was *in vitro* transcribed as described previously (Singh et al. 2014) from a DNA template ordered from Integrated DNA Technologies (IDT). See Table S8 for the DNA template primers. Embryo microinjection in C57BL/6J zygotes was performed as described previously using 50ng/uL of sgRNA and 50ng/uL of Cas9 mRNA (TriLink Biotechnologies). The resulting 11bp deletion was identified with Sanger sequencing of genomic DNA. Editing of the allele generated a novel *Bst*UI restriction site which was used to distinguish between wild-type and mutant alleles after PCR amplification (see Table S8 for genotyping primers). Generation and genotyping of *Fancm*^*em1/Jcs*^ animals was described previously (McNairn et al. 2019).

### Cell Lines [related to Figure S1]

V6.4 mESCs (You et al. 1998) were maintained under traditional mESC culture conditions (Tremml et al. 2008). Primary C57BL/6J MEFs were isolated from E13.5 embryos in which organs were removed and the remainder of the embryo was trypsinized to make a cell suspension. Cells were cultured in media comprised of DMEM with 10% FBS, 1X nonessential amino acids, and 100 units/mL penicillin-streptomycin.

### Cell Cycle Analysis

Fetal gonads from embryos whose mothers were either treated or untreated with radiation were dissected and pooled according to treatment condition and sex. Fetal gonads were disaggregated and dissociated into a single cell suspension using 0.25% trypsin-EDTA and 20μg/mL of DNase I for 15 minutes at 37°C. Trypsin was deactivated with 10% FBS. Suspensions were stained for DNA content with Hoechst 33342 (ThermoFisher 622495) and propidium iodide (PI) (Life Technologies P3566) for dead cell exclusion. Single cell suspensions were labeled for 30 minutes with 100μg Hoechst 33342 in a 33°C water bath shaking at 150rpm. Prior to cell cycle analysis samples were strained through a pre-wetted 40μm filter and labeled with propidium iodide (0.25μg/mL). Cell cycle analysis was performed using FCS Express 6 software. Statistical comparisons of cell cycle graphs were performed using GraphPad Prism8 using unpaired non-parametric Mann-Whitney tests.

### RNA-seq Sample Preparation and Gene Expression Analysis

GFP+ PGCs were purified via FACS and total RNA was isolated using Trizol-LS (Thermo Fisher) according to the manufacturer’s instructions. RNA quality was assessed by spectrophotometry (Nanodrop) to determine concentration and chemical purity (A260/230 and A260/280 ratios) and with a Fragment Analyzer (Advanced Analytical) to determine RNA integrity. Ribosomal RNA was subtracted by hybridization from total RNA samples using the RiboZero Magnetic Gold H/M/R Kit (Illumina) and the rRNA-subtracted samples were quantified with a Qubit 2.0 (RNA HS Kit; Thermo Fisher). TruSeq-barcoded RNA-seq libraries were generated with the NEBNext Ultra II RNA Library Prep Kit (New England Biolabs) and each library was quantified via Qubit 2.0 (dsDNA HS kit; Thermo Fisher) prior to pooling. For analysis, reads were trimmed to remove adaptor sequences and low quality reads using Cutadapt v1.8 with parameters: -m 50 –q 20 –a AGATCGGAAGAGCACACGTCTGAACTCCAG –match-readwildcards. Reads were then mapped to the mm10 mouse reference genome/transcriptome using Tophat v2.1. For gene expression analysis, Cufflinks v2.2 (cuffnorm/cuffdiff) was used to generate FPKM values and statistical analysis of differential gene expression (Trapnell et al. 2010)

Gene Ontology analyses were conducted using PANTHER Classification System (Mi et al. 2019) and heatmaps were generated using heatmapper.ca (Babicki et al. 2016)

### Transposable Element Expression Analysis

To conduct transposable element differential expression analysis, the software package TEtranscripts (Jin et al. 2015) was used with the default settings and the associated mm10 TE annotation GTF file.

### Small RNA-seq Sample Preparation and Analysis

Total RNA was isolated as described above and the presence of small RNAs (smaller than 200 nucleotide fragments) was detected with a Fragment Analyzer (Advanced Analytical). TrueSeq-barcoded RNA-seq libraries were generated with the NEBNext Small RNA Library Prep Kit (New England Biolabs) and size selected for insert sizes ∼18-50bp. Each library was quantified with a Qubit 2.0 (dsDNA HS Kit; Thermo Fisher). For piRNA analysis, reads were trimmed using Trim Galore! and then run through piPipes small RNA-seq pipeline with alignment to the mm10 reference genome (Han et al. 2015) and using the default settings.

### Fertility Tests, Sperm Counts, and Testis Histology

Methods were conducted as described in (Bloom and Schimenti 2020). Statistical comparisons were performed using GraphPad Prism8 using unpaired non-parametric Mann-Whitney tests.

### Single Cell DNA-sequencing and Analysis

Round spermatids were isolated from mice of the following genotypes: wild-type, *p21*^*-/-*^, *Fancm*^*-/-*^, *Fancm*^*-/-*^ *p21*^*-/-*^ at postnatal day 26 using fluorescence activated cell sorting (FACS). To FACS spermatids, Vybrant DyeCycle Violet Stain was used to label cellular DNA according to the manufacturer’s instructions (ThermoFisher Scientific V35003) and PI was used to exclude dead cells as described in the Cell Cycle Analysis methods section. Spermatids were individually sorted into PCR tubes and flash frozen prior to DNA amplification.

Single cells were subjected to AccuSomatic single-cell multiple displacement amplification (Dong et al. 2017; Milholland et al. 2017) for whole-genome sequencing by Singulomics, New York, NY. Sequencing libraries were also prepared by Singulomics. Amplicons were prepared using the NEBNext^®^ DNA Library Prep Kit following manufacturer’s recommendations. The libraries were analyzed for size distribution by an Agilent 2100 Bioanalyzer and quantified using real-time PCR. The libraries were pooled according to their effective concentrations and sequenced on Illumina NovaSeq6000 sequencer with 150 bp paired-end model using the NovaSeq6000 SP Reagent Kit. Approximately 1 Gb of sequencing data was generated per cell. Parental genomic DNA was isolated from spleens and subjected to 100 bp paired-end whole genome sequencing using BGI’s DNBseq (BGISEQ-500) platform.

Samples were aligned to the mm10 reference genome using BWA-MEM 0.7.17 (Li 2013) and variants were called from sorted BAM files using Platypus 0.8.1 (Rimmer et al. 2014). The following Platypus settings were applied to all samples: --assemble=1 –assemblyRegionSize=5000 –maxSize=5000. BCFtools 1.9 was used to filter out variants present in both individual germ cell genomes and parental genomes in order to identify germ cell-specific variants. Statistical significance was assessed using the Kruskal-Wallis test controlling for multiple comparisons using a Benjamini False Discovery Rate (FDR) correction. Genome browser images were generated using the Integrative Genomics Viewer (IGV) (Robinson et al. 2011).

## Data Availability

Raw data files from the RNA-seq, small RNA-seq and scDNA-seq experiments have been deposited onto the GEO database with accession number (pending ms review; available upon request for reviewers).

## Acknowledgments

This work was supported by National Institutes of Health grants T32HD057854 to J.C.B. and R01HD082568 to J.C.S. The authors would like to thank J. Grenier and Cornell’s Transcriptional Regulation and Expression Facility for assistance with RNA-sequencing and small RNA-sequencing experiments (P50-HD076210) and J. Glaubitz for helpful discussions regarding single cell DNA-sequencing analysis. R. Munroe and C. Abratte of Cornell’s Stem Cell and Transgenic Core Facility generated the *Fancm*^*em1/Jcs*^ and *p21*^*em1/Jcs*^ mouse lines with partial support from the Empire State Stem Cell Fund (contract # C024174).

## Author Contributions

J.C.B. conducted the experiments described and performed data analysis. J.C.S supervised all aspects of the work. J.C.B. and J.C.S wrote the manuscript.

